# Cooperation maximizes biodiversity

**DOI:** 10.1101/2024.10.22.619656

**Authors:** Oscar Godoy, Fernando Soler-Toscano, José R. Portillo, Antonio Suárez, José A. Langa

## Abstract

Cooperation, the mutual benefit that individuals of different species obtain when they interact together, is ubiquitous in nature. Despite their importance, most of all current ecological theories have been formalized focusing on negative interactions such as competition or predation. The role of cooperation, or other types of positive interactions including facilitation and mutualism, has not been fully addressed, or, if so, always in combination with negative interactions. This fact limits our understanding of the unique features by which cooperation as opposed to competition promotes biodiversity. To address this gap, we introduce here cooperation into structural stability, a general framework to understand how species interactions and environmental variability determine the long-term persistence of species within communities. Compared to a pure competitive case, cooperation promotes three distinctive features. First, cooperation increases the opportunities for species to coexist. This feature increases the persistence of species with contrasted phylogenetic, functional, and demographic strategies that the environment would otherwise filter. Second, cooperation creates intertwined biodiversity where the existence of some species begets the presence of others. Third, cooperation promotes multistability by changing the dynamics of community assembly due to variations in environmental conditions. In conclusion, we present a fully operational framework to understand the unique ecological roles of cooperation in nature. It indicates that cooperation as opposed to competition maximizes the maintenance of biodiversity.

## 1 Introduction

Ecologists have extensively studied biotic interactions because of their fundamental importance for the maintenance of biodiversity. However, most of the major past and current developments focus on the study of negative interactions such as competition, predation, or parasitism ([1, 2, 3, 4, 5, 6]). Cooperation, the mutual benefit that individuals of different species obtain when they interact together, as well as other types of positive interactions such as facilitation or mutualism, are not yet included in general principles of ecology and evolutionary biology despite numerous attempts ([7, 8, 9, 10, 11, 12]). Some researchers argue that cooperation plays a secondary role due to a combination of historical aspects associated with the Darwinian struggle for existence and the economic growth based on capitalism [13]. However, we advocate that this bias has not yet been overcome for two main reasons. On the theoretical side, conceptual developments and associated mathematical attempts have not elucidated the distinctive features by which cooperation, as opposed to competition, promotes biodiversity (but see some progress in 14). This is mostly because cooperation and other sources of positive effects are considered in combination with negative interactions rather than on their own. In such a combination, cooperation is often seen as a buffering mechanism against the negative effects of interactions and environmental conditions on biodiversity. For example, it is assumed that the obligate cooperation of insects acting as pollinators or fungi establishing their mycorrhizae promotes diversity because it alleviates the strong competition between their plant counterparts ([15, 16, 17, 18]). Likewise, neighbors increase species’ performance at low density by improving local climatic conditions (e.g. [19]). On the empirical side, studies narrow the prevalence of cooperation in stressful habitats such as arid/desert ecosystems, marshlands, or alpine/cold environments, in which abiotic conditions do not allow species to thrive ([20, 21, 22]). However, there is now enough cumulative evidence that cooperation shapes both evolutionary and ecological processes across climatic conditions, ecosystem types, and kingdoms of life in obligate and facultative forms ([7, 23, 24, 25, 26]).

This perspective seeks to fill a significant gap in ecological understanding by exploring which are the distinctive features that cooperation, as opposed to competition, promotes in the assembly of ecological communities and the maintenance of biodiversity. To address this gap, we integrate cooperation into general principles of ecology by using the framework of structural stability. In the following sections, we first briefly explain the main principles from the field of structural stability (Box 1). Then, we illustrate how coupling them with cooperation provides three distinctive features that make cooperation as important or even more important than competition. This is because of the capacity of cooperation to maximize biodiversity, resilience, and multistability of ecological systems.

## 2 Cooperation enlarges the opportunities for species to coexist

Competition for resources or the share of natural enemies are often invoked as key forces shaping the diversity and composition of ecological communities. Different ecological theories posit that when a community is dominated by competition, the range of opportunities for species to coexist depends on the ratio of intraspecific versus interspecific interactions, that is, how species limit themselves compared to how they limit others. The main prediction is that the stronger intraspecific exceeds interspecific competition, the more likely species coexist despite marked differences in performance ([27, 28]). This is because it is assumed that species differ more in the resources they use and the natural enemies they share, and such niche differences act as stabilizing mechanisms limiting species when dominant but buffering them from extinction when rare ([29, 30]). Eventually, the relative strength of competitive interactions combined with their intrinsic performance defines two clear competitive outcomes (Fig. 2a). Species are predicted to coexist (i.e., long-term persistence) when the ratio of intra versus interspecific competition, technically called the feasibility domain, can accommodate differences in species’ performance. Conversely, competitive exclusion of the inferior competitor occurs when the ratio between intra and interspecific competition cannot accommodate such differences in species performance. There is a third possibility, called environmental filtering, in which none of the species can be present in the community because the environment is too harsh for the species to thrive.

**Figure 1:**
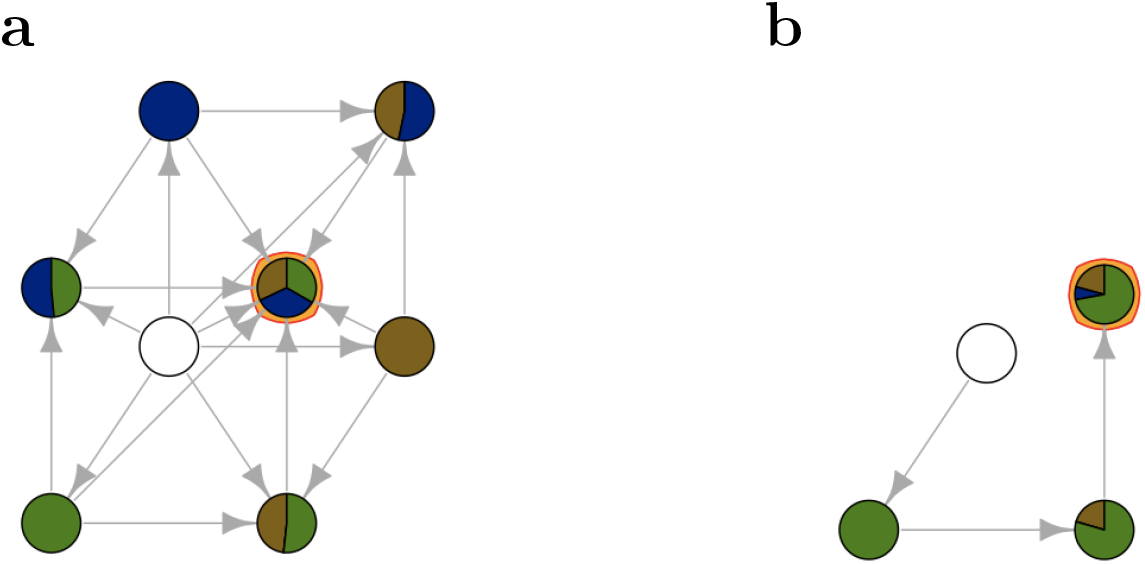
Informational structures (ISs) of a three-dimensional cooperative Lotka-Volterra model. Different ISs emerge when cooperation is combining positive intrinsic growth rates for all species (**a**, *r*_1_ = 0.252(brown), *r*_2_ = 0.322(blue), *r*_1_ = 0.287)(green)) or some (**b**, *r*_1_ = −0.022, *r*_2_ = −0.155, *r*_1_ = 0.475, same color order). In **a**, the IS contains all possible stable subcommunities (circles), including the empty community (white circle) and their connections. Therefore, it is possible to end in the community with maximum richness by departing from any species combination. In **b**, the IS shows an extreme case of intertwined biodiversity. The community of three species only emerges, when the presence of species 3 (green) favors the persistence of species 1 (brown), which in turn cooperates with species 2 (blue). Note that the relative abundance of each species in the community is proportional to its slice size in the pie chart. This comparison of **a** and **b** shows that both communities sustain the same biodiversity while exhibiting completely distinct interdependencies, precisely expressed by the underlying structure.

**Figure 2:**
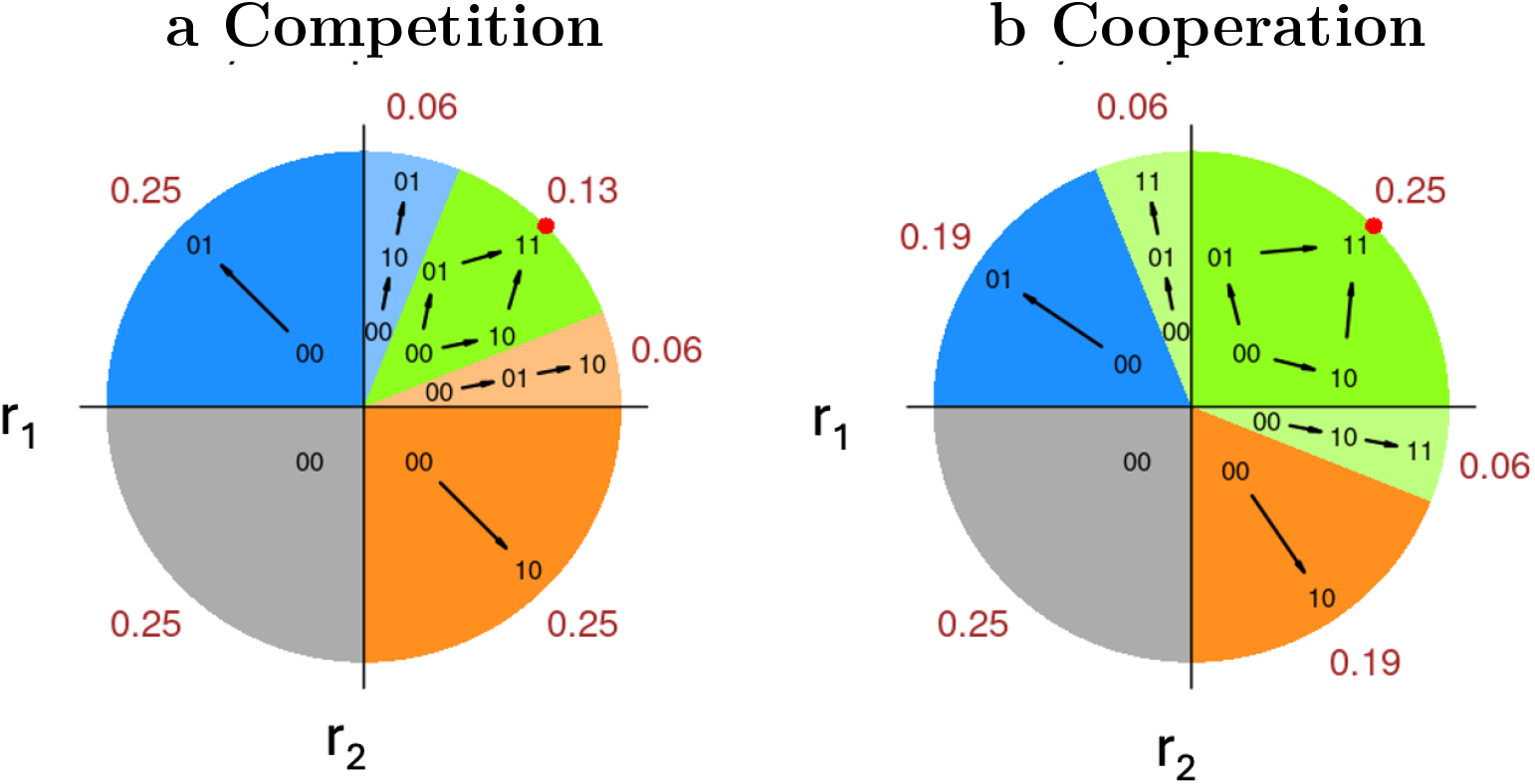
A structural stability framework to understand the maintenance of biodiversity. The range of variation in intrinsic growth rates of species (*r*_1_,*r*_2_) for a given network of interactions defines distinctive outcomes and their assembly dynamics. These outcomes are represented by different colored areas (red numbers represent the proportion of each area). **a** For competition, the green area represents the range of species’ intrinsic growth rates compatible with species coexistence. That is, departing from an empty community, it is viable and stable to end in a community composed of both species (1,1). The light blue and orange areas represent cases where the inferior competitor is excluded, communities either as (1,0) or (0,1). The dark orange and blue areas correspond to cases in which species cannot cope with environmental conditions and therefore are filtered by the environment, and, finally, the gray area indicates the case in which none of the species can persist (0,0). **b** Under cooperation, three unique processes are observed. First, the green area is enlarged, indicating that species can coexist under a wider range of environmental conditions. Second, both regions of competitive exclusion disappear, and instead, a new region emerges (light green). We call this area the case of “intertwined diversity” in which the presence of one species helps another to persist thanks to cooperation. Third, in cooperation, different community assembly structures that end in the maximum level of biodiversity (1,1) arise, promoting the multi-stability of the ecosystem. See Fig. 3 for an easier interpretation of the link between cooperation and multistability. For illustration, parameterization of the LV model was performed according to symmetric interspecific effects between species, Competition: (*r*_1_=*r*_2_= 1, *α*_12_=*α*_21_= -0.4, *α*_11_=*α*_22_= -1). Cooperation: (*r*_1_=*r*_2_= 1, *α*_12_=*α*_21_= 0.4, *α*_11_=*α*_22_= -1)

Under this general framework, we observe marked differences when cooperation replaces competition. When species are involved in interspecific positive interactions while maintaining self-limiting effects, it is straightforward to observe an increase in the opportunities for species to coexist. In technical terms, the feasibility domain is larger (Fig. 2b), and the more cooperation in the system, the larger the feasibility domain regardless of the number of species (see Fig. 3 for an example with three species). This increase, which was previously observed in conceptual ([8]) and mathematical ([31]) work, has a main and so far overlooked implication: communities under cooperation can withstand greater asymmetries in species’ performance without losing species. One end of such asymmetries includes a subset of species in a community showing negative intrinsic growth rates, indicating that species cannot cope with local environmental conditions by themselves in the absence of interactions. This is likely why cooperation has historically been linked to unfavorable environments for plant growth through the stress gradient hypothesis (e.g., [32]). Although we do not deny that cooperation (also called facilitation in the plant literature) can be more common in these environments, and it has greater effects on maintaining diversity ([33]), we believe that restricting cooperation to particular environmental conditions limits our perception of the ubiquity and importance of cooperation in nature.

**Figure 3:**
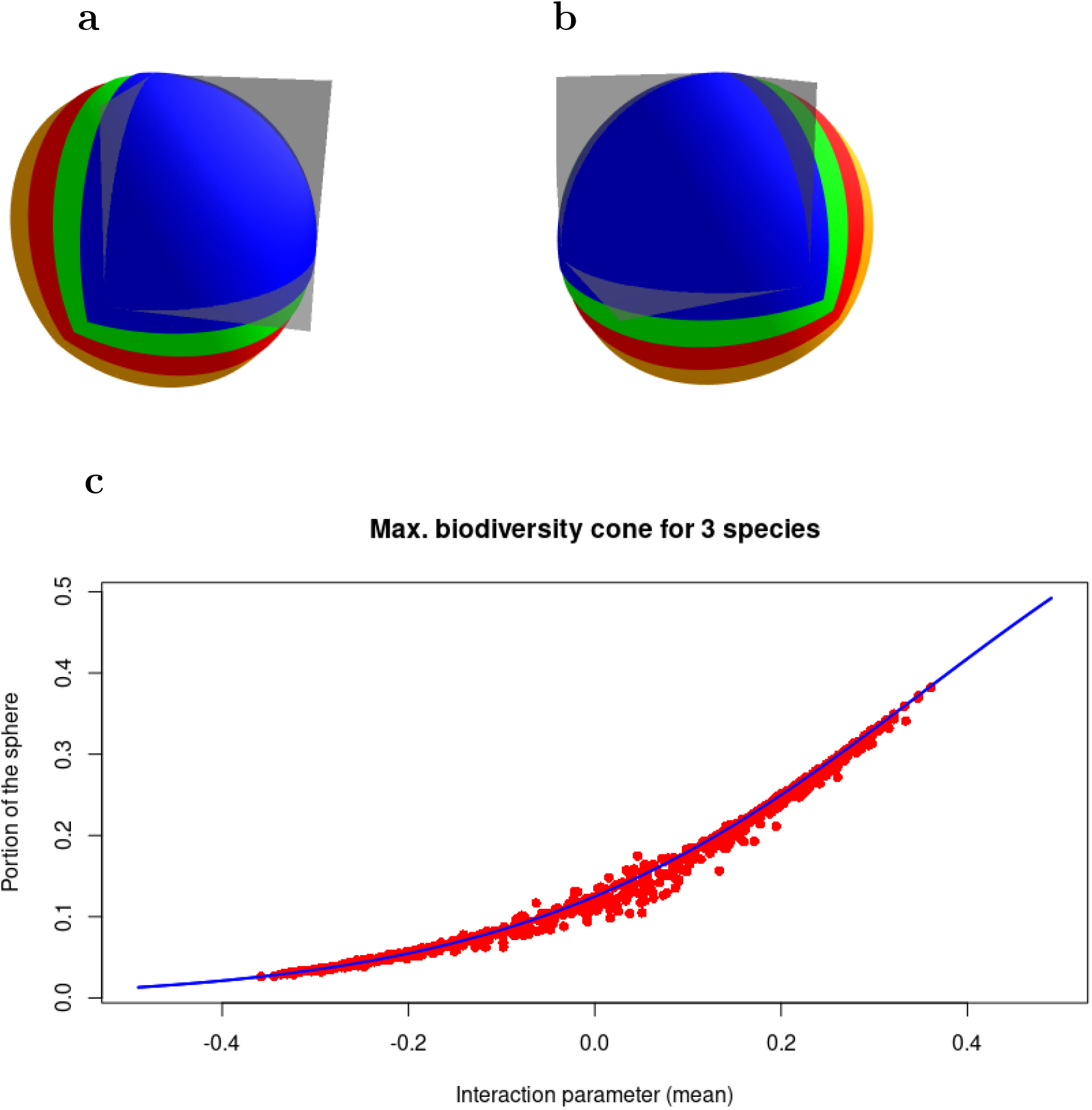
Increase in the size of the cone of maximal diversity with cooperation. **a** (left side) and **b** (right side) illustrate the effect that an increase in the prevalence of cooperation has on enlarging the cone of maximum diversity. This cone, technically called the feasibility domain, contains the range of intrinsic growth rates compatible with the persistence of all three species. More specifically, the color change is associated with an increase in the cone size for the following interspecific values (blue *α*_*ij*_= 0.1, green= 0.2, red= 0.3, orange= 0.4, intraspecific values are held constant at *α*_*ii*_ = *α*_*jj*_ =-1). **c** Changes in the portion of the sphere covered by the cone of maximum diversity (y-axis) as a continuous function of changes from competition to cooperation (x-axis). Competition are negative interaction values *α*_*ij*_*<*0) and cooperation are positive interaction values *α*_*ij*_*>*0). In this relationship, the blue line corresponds to a matrix with *α*_*ii*_ = −1 and constant *α*_*ij*_ for *i* ≠ *j*. Red dots correspond to matrices with *α*_*ii*_ = −1 and random *α*_*ij*_ for *i* ≠ *j*. The mean value of the *α*_*ij*_ is considered. Note that the relationship is 1) highly-dependent on the total sum of *α*_*ij*_ coefficients, and 2) always positive and non-saturating, which illustrates that the more cooperation the larger the opportunities for species to coexist, and always greater than under competition.

Other cases in which cooperation also allows to withstanding large asymmetries in species’ performance arise from disparate demographic strategies. Earlier ecological literature did not consider this possibility because it was focused more on understanding how closely related species coexist under competition ([4]). Because competition reduces the opportunities for species to coexist, predictions were that they could do so only when their fitness tends to be similar ([4, 29]), something frequently observed among close relatives. However, by including cooperation, we can expand our question to ask how distantly related species can coexist. When species with disparate functional and demographic characteristics interact, it is likely to observe slow-versus fast-growing strategies. These strategies, which are evolutionarily conserved, make species capitalize either on high offspring production and fast reproduction, or on slow growth and survival ([34]). According to theory, contrasted growing strategies are harder to maintain under competition, yet the prevalence of cooperation in an ecological system can reverse this outcome by promoting communities with a wide variety of physiological, functional, and phylogenetic characteristics as some recent examples suggest ([10, 35]).

Another final source of large asymmetries in species’ performance can come from dispersal. It is common to observe that the community pool is composed of core species adapted to local conditions plus a set of species that have stable sources elsewhere and occasionally reach new locations by dispersal. In these new environments, dispersed species often struggle to perform well and fail to persist due to a combination of competition and inadequate adaptations to the local conditions ([36]). But thanks to enlarging the opportunities for species to coexist, cooperation acts as a mechanism that will allow the demographic growth and the persistence of these “newcomer” species while not reducing the resident pool. If the positive effect of cooperation on dispersal happens repeatedly across space, we could observe entire communities with larger species distribution across contrasted climatic and edaphic conditions. This is something that has not been extensively tested yet, but it has been suggested to occur in harsh environments as a process that enlarges the edges of species distributions ([37]), and in the successful establishment of exotic species ([38]).

## 3 Cooperation promotes intertwined biodiversity

A second important aspect is that cooperation reduces species losses when competition would otherwise suppress the inferior competitors. This is because competitive exclusion is no longer an outcome of ecological interactions (Fig. 2a). Therefore, cooperation tends to maximize the persistence of the whole species pool of the community. Instead of competitive exclusion, a new outcome emerges at the edges of the feasibility domain (Fig. 2b). In this new area that we called “intertwined diversity”, coexistence is the outcome of cooperation. That is because species that otherwise cannot persist by themselves, they can now do so thanks to the presence of other cooperative species. Thanks to this striking phenomenon, in which the presence of the second species is contingent upon the presence of the first, an ecological community can maintain the same maximum levels of diversity as in the competitive case. So, cooperation should not be viewed as a process promoting the persistence of a few species in stressful environments. Rather, cooperation can promote high levels of species richness, and in fact, the more the number of species, the more likely these interconnections to occur by different assembly paths, thereby facilitating the coexistence of all species through mutual support (Fig. 4).

**Figure 4:**
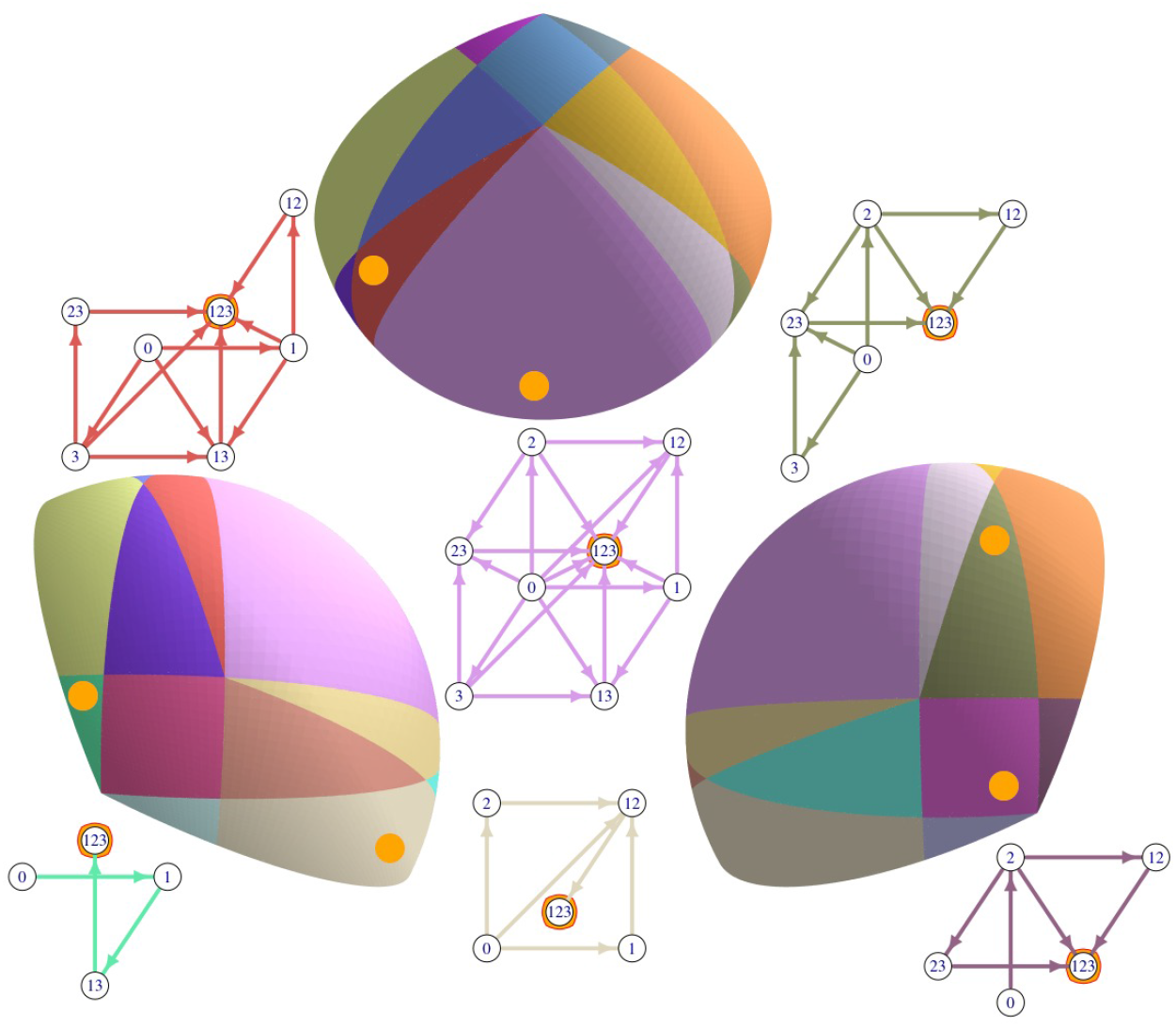
Assembly structures for communities of three species under cooperation following Lotka-Volterra dynamics. The same assembly structure can be viewed from the front, left, and right sides. As illustrated in (Fig.2) for two species, cooperation enriches the cone of maximal biodiversity by promoting new opportunities for species to coexist under the mechanism of intertwined biodiversity. However, while the number of new regions is two for a community composed of two species, the number of regions explodes to 22 when considering three species. Thereby, higher species richness rapidly promotes multistability thanks to the interconnections between stable communities, and species coexistence becomes more intricate through mutual support. Still, the region that emerges under competition (see purple region) is maintained when cooperation operates. However, the rest of the novel regions appear outside the positive octant, which indicates that the mechanisms of intertwined diversity are at play although with different strengths across assembly structures. Particularly noteworthy is the region where there is a single path to assemble the community (see green region). This situation occurs when cooperation combines with the case of all species except one showing negative intrinsic growth rates. Although for four or more species such processes of multistability and intertwined diversity can not be illustrated, the same phenomena of intertwined diversity and multistability can be observed for an arbitrary number of species.

The result in which the main cause of diversity is diversity itself has been proposed in other ecological and evolutionary contexts with contrasted mechanisms at play. For instance, historical contingencies in community assembly such as order and/or timing of arrival can change the outcome of interspecific interactions ([39]). Under such priority effects, some species can show positive population growth if early colonizers modify local environmental characteristics that favor settlement and growth of a second species ([40]). This cooperative priority effect differs from cooperation itself, as the former depends on the order of events. Meanwhile, co-operation as we present it here is a deterministic process that consistently maximizes diversity regardless of which species is present first in the system. Another situation in which diversity begets diversity has been proposed within an evolutionary context. As new species form, they create novel opportunities for others to exploit and feed on them ([41]). However, under the case of intertwined diversity, it does not arise from the combination of positive and negative interspecific interactions found in food webs but rather only from positive ones.

Because intertwined diversity occurs when each species supports others regardless of their identity, the network of interactions sustaining the persistence of species constitutes a cohesive entity wherein individual components would not thrive in isolation, forming an interdependent whole. In this case, such interdependence is of paramount importance for community resilience to withstand environmental perturbations. When indirect competitive interactions rather than cooperation maintain biodiversity, a loss of one species can concomitantly produce the extinction of others ([42]). These cascading effects are common in communities ensemble by a combination of negative and positive interactions ([43, 44]). Under cooperation the loss of biodiversity due to cascade extinctions can range from negligible effects when the species extinct presented strong negative growth rates to catastrophic effects when the species that goes extinct is the only one that can thrive alone and support the rest of the species (Fig. 4). Therefore, cooperation can produce extinction cascades, but it remove them depending on the species’ demographic characteristics.

Finally, the positive effects of cooperation on biodiversity are aligned with work done in other disciplines that might be considered to be unrelated to biological sciences. In neuroscience, for example, cooperation among different regions of the human brain has been explored to successfully distinguish between several states of consciousness, sometimes in patients with severe alterations affecting brain dynamics ([45, 46]). Higher levels of consciousness are associated with interlinked nodes, thus, such cooperation in the human brain represents a measure of health. This finding opens the question whether the degree of cooperation can be considered a measure of ”ecosystem health” due to its positive effects on biodiversity.

## 4 Cooperation promotes multistability

The final novel aspect of cooperation is that it promotes multistability in the following manner. Competition produces only a single structure of community assembly (see Fig. 4, purple region) generally called Informational Structures (ISs) [47, 48] or assembly structures/invasion graphs in ecology ([49, 50]). This assembly structure is composed of multiple stable communities and the connections between these subcommunities, which overall define the different paths by which a set of species can be assembled (Box 1). However, the situation is dramatically different under cooperation. Cooperation promotes multiple of these community assembly structures. These new cooperative structures, where intertwined diversity occurs, emerge around the competitive case, resembling petals radiating from the center of a flower (Fig. 4). This bifurcation of assembly structures implies multistability as different transient and asymptotic dynamics for the community are expected when species vary their intrinsic growth rates. The total number of these structures of community assembly *s*_*n*_, each one with its topology, depends on the number of species *n* considered in the community. There is a total of 6 assembly structures for two species (Fig. 2b) and the number increases up to 34 for three species (Fig. 4). For four species, although it can not be illustrated the total number of assembly structures is 672, and the sequence continues at 199572, and 12884849614 for five and six species, respectively. Beyond six species, which can be considered a system with low-medium diversity, the number of new assembly structures is currently unknown due to computational limitations [51]. More specifically, this sequence of numbers (labelled as *A*224913 in the On-Line Encyclopedia of Integer Sequences [52]) follow this formula 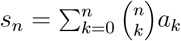 where *a*_*k*_ is the number of antimatroids (assembly structures) with *n* labelled items (species). Antimatroids were discovered almost a century ago in mathematics [53]. They present numerous properties that have been applied to diverse fields such as game theory [54] and economics [55], but they have never been introduced into ecology and evolution. Here, we show the cooperative structure of community assembly are in fact antimatroids, and they hold an enormous power to promote multistability.

The study of how an ecosystem can transition from one state to another due to external or internal forces with consequences for the structure and function of the ecosystem has been a wide subject of research in ecology for the past 50 years ([56, 57, 58]). Perhaps the most illustrative examples of multistability are those involved in the early study of alternative stable states in lakes between a diverse and productive ecosystem and another impoverished ([59, 60]). More recently, similar concepts have been applied to study shifts between semi-arid and desert ecosystems ([61, 62]). Yet, in all these efforts the link between alternative stable states and positive interactions has not been established solidly ([63, 64]). Including cooperation in structural stability can fructify this connection because it provides insights into two important aspects of multistability not previously considered. The first aspect is that multistability under cooperation is likely a common phenomenon. The main argument to make this hypothesis is that these new regions of community assembly are small (compare them to the size of the purple region that is the only one that emerges under competition) (Fig.4), and hence, temporal changes in species’ intrinsic growth rates, which are associated with temporal changes in environmental conditions, will continuously move the community between assembly structures ([65]). Therefore, different communities varying in species richness, composition, and abundance will be promoted over time as has been recently shown with bacterial communities ([65]). The second aspect is that multistability goes beyond considering only a few alternative states that typically arise when considering a low number of species, or variables that summarize community properties such as biomass or productivity ([66]). Recent work has shown that a plethora of multiple stable states (termed “cliques”) emerge in scenarios of high diversity ([67]). In the structural stability framework, these cliques are precisely the different regions of assembly we observe under cooperation (Fig.4). Although these cliques can be currently quantified, it is unclear how they are connected. Here, by mapping the position, and the extent of the different assembly structures, we can now understand such connections between cliques. This means that we can predict the more likely transition between cliques, and therefore between different stable states, depending on the magnitude and direction of the environmental change.

## 5 Cooperation and competition mix

In previous sections, we have explored the three distinctive features by which cooperation as opposed to competition promotes biodiversity. However, the most likely scenario in nature is that cooperation and competition mix. This is something pervasive in communities with obligated forms of cooperation such as mutualistic systems ([16, 68]). It is also frequently documented in plant communities where cooperation is facultative and it can change to competition depending on environmental conditions ([25, 69]). Indeed, this cooperation can be asymmetric in many different ways. A species cooperates with many others but not the other way around. This is a well-known case of engineering species or nurse plants ([70, 71]). It is also common to observe a pair of species cooperating while others compete. This second case describes the interaction between a plant and a pollinator, and between two plants, respectively ([72]). And finally, a species can cooperate with another while competing with a third one. This phenomenon is common in ecological communities considering a single trophic level such as plant or plankton communities ([25, 73]).

When cooperation and competition act together, the three features unique to cooperation still hold. However, the benefits are not now for all species within the community but for particular species. In other words, some species will become dominant because they benefit more from withstanding larger variations in environmental conditions. Some species will be only present in the community due to the presence of others, and lastly, communities will vary their composition between different stable states depending on their ability to grow. Assessing all these nuances is complex but fortunately, it is now possible with recent tools developed under the framework of structural stability ([74]).

## 6 Future directions and conclusion

With the inclusion of cooperation into a general framework of species coexistence and community assembly many future directions of research open. Among the most pressing issues to address is to determine empirically how these three distinctive features of cooperation are realized across ecosystems. In this quest, we believe is key to understanding the limits to cooperation between species. While in some systems species may cooperate or not depending on external factors such as climate, others are forced to cooperate if they do not want to become extinct. Moreover, the connections between community ecology and macroecology through species interactions are on the rise ([75]), and it is a good opportunity to quantify rigorously the importance of cooperation for species and community distributions across large spatial extents rather than their traditional restriction to harsh environments. Taken together, we show here that cooperation maximizes biodiversity by increasing the functional strategies, resilience, and multistability of ecological communities.

### Box 1 — Structural stability in a nutshell

Structural stability measures the range of variation in species performance, that is, variation in intrinsic growth rates, among interacting species compatible with their long-term persistence. The larger is this range, technically called feasibility domain, the greater can be differences in performance (Fig. 2). Therefore, an ecological community is structurally stable when variation in model parameters, in particular in species’ intrinsic growth, affect species abundances but no species go extinct. There are indeed many models to understand and predict temporal changes in species abundances, but a model widespread used in ecology and central to the use of structural stability is the linear Lotka-Volterra (LV) model, which represents a balance between tractability and complexity, and can generate population dynamics with the range of complexity of more complex models [76, 77, 6].

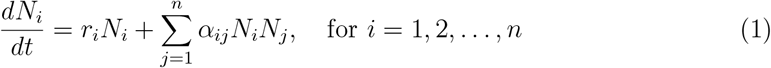

In this system of equations:

- *N*_*i*_ denotes the population size of species *i*.
- 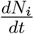 represents the rate of change of the population size of species *i* over time.
- *r*_*i*_ represents the intrinsic growth rate of species *i* in the absence of interactions.
- and *α*_*ij*_ refers to the interaction coefficient between species *i* and species *j*.

The Lotka-Volterra equations capture two key ecological processes. On the one hand, the term *r*_*i*_*N*_*i*_ describes the intrinsic growth of species *i* in the absence of interactions with other species, reflecting the species’ capacity to reproduce and increase its population size. This term is negative when species cannot cope with local environmental conditions alone (*r*_*i*_ *<* 0), and therefore, are filtered by the environment [78]. On the other hand, the term 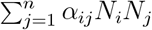 accounts for the interaction strength and sign between species within the community in a phenomenological way. Negative values of *α*_*ij*_ indicate competition whereas positive values indicate cooperation. That is, competitive interactions mean that the population growth of one species will be reduced by the presence of others while cooperative interactions mean the opposite.

The structural stability tells us not only the range of intrinsic growth rates compatible with the persistence of interacting species but also the community assembly structure that the system will follow for different combinations of model parameters. Therefore, such a structure, technically called Informational Structure (IS), connects several fields of biology such as species coexistence, community ecology, global change, conservation biology, and restoration ecology. More specifically, an IS describes the dynamics of interacting species according to a population model from which emerges the skeleton of a global attractor according to two elements. On the one hand, it contains the set of subcommunities (i.e., communities that do not contain all species pool) that are stable, and on the other hand, it also includes all the relationships between these subcommunities. Therefore, an ISs generates a network with explanatory power for the dynamics corresponding to any initial state of the community, which includes all possibilities of species assembly as well as extinction cascades (Fig. 1, Box). Describing an IS is still a complex subject of research in mathematics ([79, 80, 50]), yet they can be fully defined when the interaction matrix of the Lotka-Volterra model is Volterra-Lyapunov stable ([81]). That occurs for instance when the strength of intraspecific interactions is greater than the sum of interspecific interactions (i.e. it is a diagonally dominant matrix). For illustration purposes, we will focus here on these types of matrix parameterizations.

